# Surveying the landscape of tRNA modifications by combining tRNA sequencing and RNA mass spectrometry

**DOI:** 10.1101/723049

**Authors:** Satoshi Kimura, Peter C. Dedon, Matthew K. Waldor

**Affiliations:** Division of Infectious Diseases, Brigham and Women’s Hospital; Department of Microbiology, Harvard Medical School; Howard Hughes Medical Institute; Department of Biological Engineering, Massachusetts Institution of Technology; Singapore-MIT Alliance for Research and Technology Antimicrobial Resistance Interdisciplinary Research Group

## Abstract

Chemical modification of the nucleosides that comprise tRNAs are diverse^1-3^. Such modifications impact tRNA structure, stability, and mRNA decoding^3,4^. Although tRNA modifications are present in all kingdoms of life^1^, the structure, location, and extent of modifications have been systematically charted in very few organisms, in part because mapping modifications to tRNA sequences has been technically challenging. Here, we describe a new approach in which rapid prediction of modified sites through reverse transcription-derived signatures in high-throughput tRNA-sequencing (tRNA-seq) data is coupled with chemical analysis and identification of tRNA modifications through RNA mass spectrometry (tRNA-SMS). As proof of concept, we applied this method to study tRNA modification profiles in two phylogenetically close bacteria, *E. coli* and *Vibrio cholerae*. Comparative tRNA-seq enabled prediction of several *V. cholerae* modifications that are absent from *E. coli* and showed the effects of various environmental conditions on *V. cholerae* tRNA modification profiles. Through RNA mass spectrometric analyses, we showed that two of the *V. cholerae-*specific reverse transcription signatures reflected the presence of a new modification (acetylated acp^3^U (acacp^3^U)), while another results from C-to-U RNA editing, a process not described before in bacteria. By combining comparative genomics with mass spectrometry, we identified a putative N- acetyltransferase required for acacp^3^U acetylation. These findings demonstrate the utility of the tRNA-SMS approach for rapid characterization of tRNA modification profiles and environmental control of tRNA modification. Moreover, our identification of a new modified nucleoside and RNA editing process suggests that there are many tRNA modifications awaiting discovery.

---

Several methods have been developed to predict the presence of modified sites from deep sequencing data of tRNAs^5,6^. To verify that deep sequencing of tRNAs (tRNA-seq) can predict the presence of tRNA modifications through detection of misincorporated nucleosides or premature termination of reverse transcription, we conducted deep-sequencing of purified total tRNA from *E. coli*, where the sites and chemical nature of tRNA modifications have been elucidated^7^. The sequencing library was constructed using a modified version of a published protocol^6^ (Supplementary Fig. 1). Reads were mapped to reference *E. coli* tRNA genes and pile up files for each locus were generated (e.g., Supplementary Fig. 2A). As expected, we observed a drop in mapped read depth and/or incorporation of mismatched bases at known modified residues (e.g., s^4^U at position 8, D at 16 and 17, mnm^5^s^2^U at 34 and acp^3^U at 47; Supplementary Fig. 2A). Heatmaps depicting the frequency of misincorporation and ratio of termination throughout all tRNAs were generated (Supplementary Fig. 2B). In total, more than half of known modifications were detected in the sequencing of *E. coli* tRNA (16 out of 28) (Supplementary Fig. 2B, Supplementary Table 1, Supplementary Data 1, and see Methods). Thus, RT-derived signatures can be used to profile many tRNA modifications.

Next, we applied tRNA-seq to define tRNA modification profiles in an organism where tRNA modifications have not been characterized: *Vibrio cholerae*, the cholera pathogen. Analysis of tRNA-seq data from stationary phase *V. cholerae* samples yielded heatmaps of misincorporation and termination similar to those of *E. coli* (Fig. 1). This observation is consistent with the conservation of most *E. coli* tRNA modification enzymes in *V. cholerae* (Supplementary Data 2). The identity of modifications introduced by *thiI, ttcA*, and *miaA* (s^4^U, s^2^C, and ms^2^io^6^A, respectively) was confirmed by analyzing tRNA from strains lacking these enzymes (Supplementary Fig. 3A-C).

**Figure 1.**
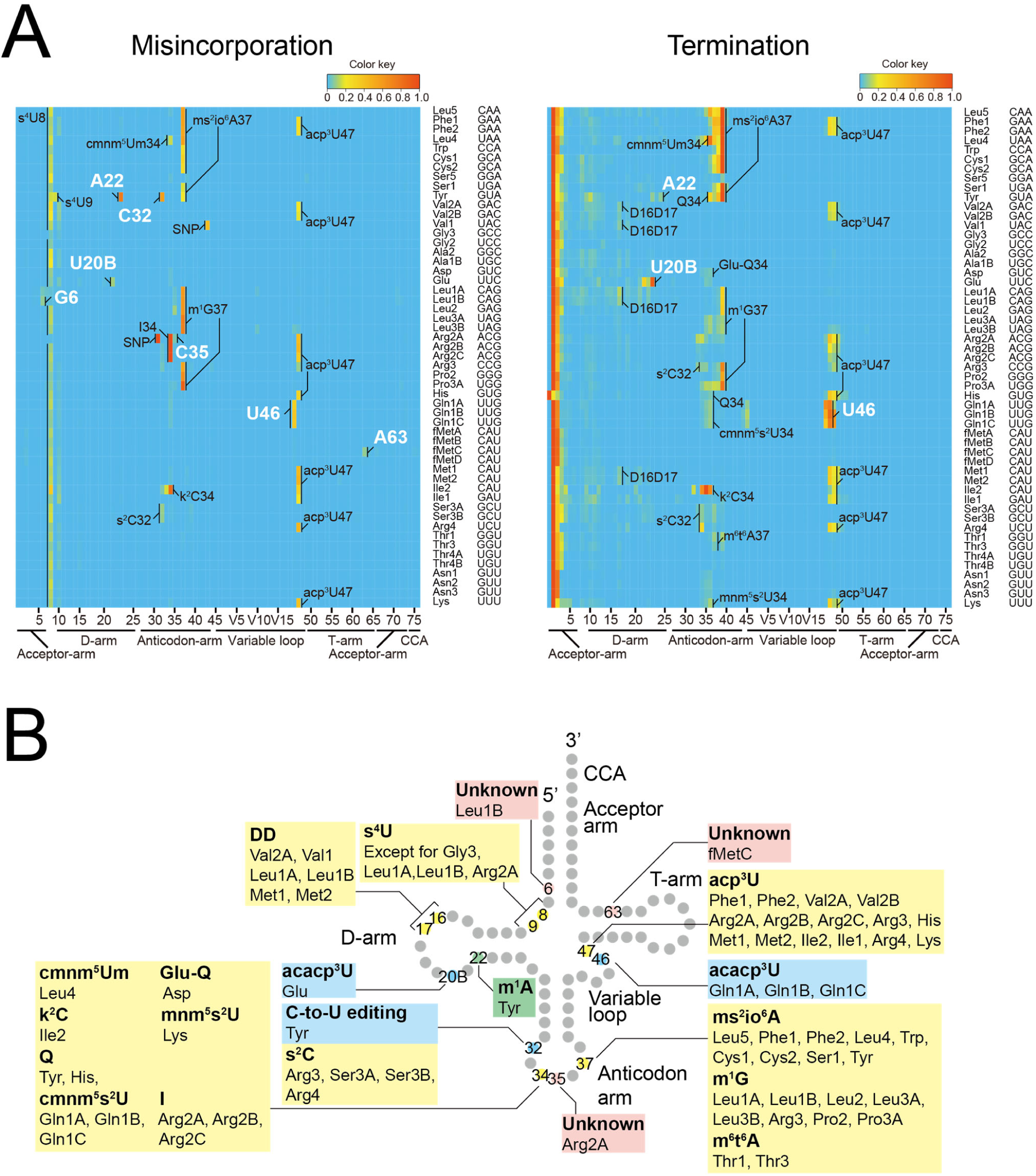
Profiling tRNA modifications in *V. cholerae* through tRNA-seq. (A) Heatmaps of frequency of misincorporation (Left) and termination of reverse transcription (Right) in a representative tRNA sample isolated from stationary phase *V. cholerae* (total of three independent samples). Positions of modifications bearing greater than 5 % of misincorporation or termination frequency are shown; the identity of presumably shared modifications with *E. coli* are indicated in black and the *V. cholerae*-specific signals are in white. Single nucleotide polymorphisms (SNPs) shown are based on whole genome sequence of C6706^18^ (Supplementary Fig. 11) are also indicated in black. (B) Schematic secondary structure of *V. cholerae* tRNAs showing sites of predicted tRNA modifications deciphered from tRNA-seq data in (A). The positions and tRNA species in which the RT-derived signatures are commonly observed in *E. coli* are shown in yellow. The positions and tRNA species that have *V. cholerae* specific signals are colored coded as green (found in other organisms but not *E. coli*), light blue (novel modifications/or editing) or pink (unknown).

We observed several misincorporation signals in *V. cholerae* that were not present in *E. coli*, including at A22 and C32 in tRNA-Tyr, U20B in tRNA-Glu, U46 in tRNA-Gln1A, Gln1B and Gln1C, G6 in tRNA-Leu1B, C35 in tRNA-Arg2A and A63 in tRNA-fMetC (Fig. 1). U20B in tRNA-Glu and U46 in tRNA-Gln1ABC were also associated with increased termination of reverse transcription (Fig. 1). In other bacteria, e.g., *B. subtilis*, A22 is methylated to m^1^A by TrmK in a subset of tRNAs^7,8^; consequently, we explored the effect of *V. cholerae’s* TrmK homolog on misincorporation at A22 in tRNA-Tyr. A *V. cholerae trmK* deletion mutant lacked the misincorporation signal at A22 in tRNA-Tyr, suggesting that *V. cholerae* contains m^1^A in tRNA-Tyr (Supplementary Fig. 3D).

The variation of modification profiles of individual tRNAs in log and stationary phase bacteria have not been systematically characterized. Comparisons of tRNA-seq patterns of samples derived from log and stationary phase *V. cholerae* cultures (Fig. 1A and Supplementary Fig. 4A) revealed a significantly higher misincorporation frequency at position 47 in a subset of log phase tRNAs (e.g., tRNA-Met), which presumably contain acp^3^U (Fig. 2A). Mass spectrometric analysis of purified tRNA-Met fragments confirmed that this modification is present at this site and is more prevalent in the log phase sample (Fig. 2B, C and Supplementary Fig. 5). Interestingly, in *E. coli*, misincorporation frequency at position 47 was higher in stationary phase (Fig. 2D), suggesting that there are species-specific mechanisms for control of modification frequency. We also determined the tRNA-seq profiles of *V. cholerae* tRNA samples derived from the cecal fluid of infant rabbits infected with the pathogen^9^. The resulting misincorporation and termination signatures were similar to those of log phase samples (Supplementary Fig. 4B vs. 4A), including modification at position 47 (Fig. 2A). Thus, tRNA-seq enables assessment of tRNA modification profiles under a variety of conditions.

**Figure 2.**
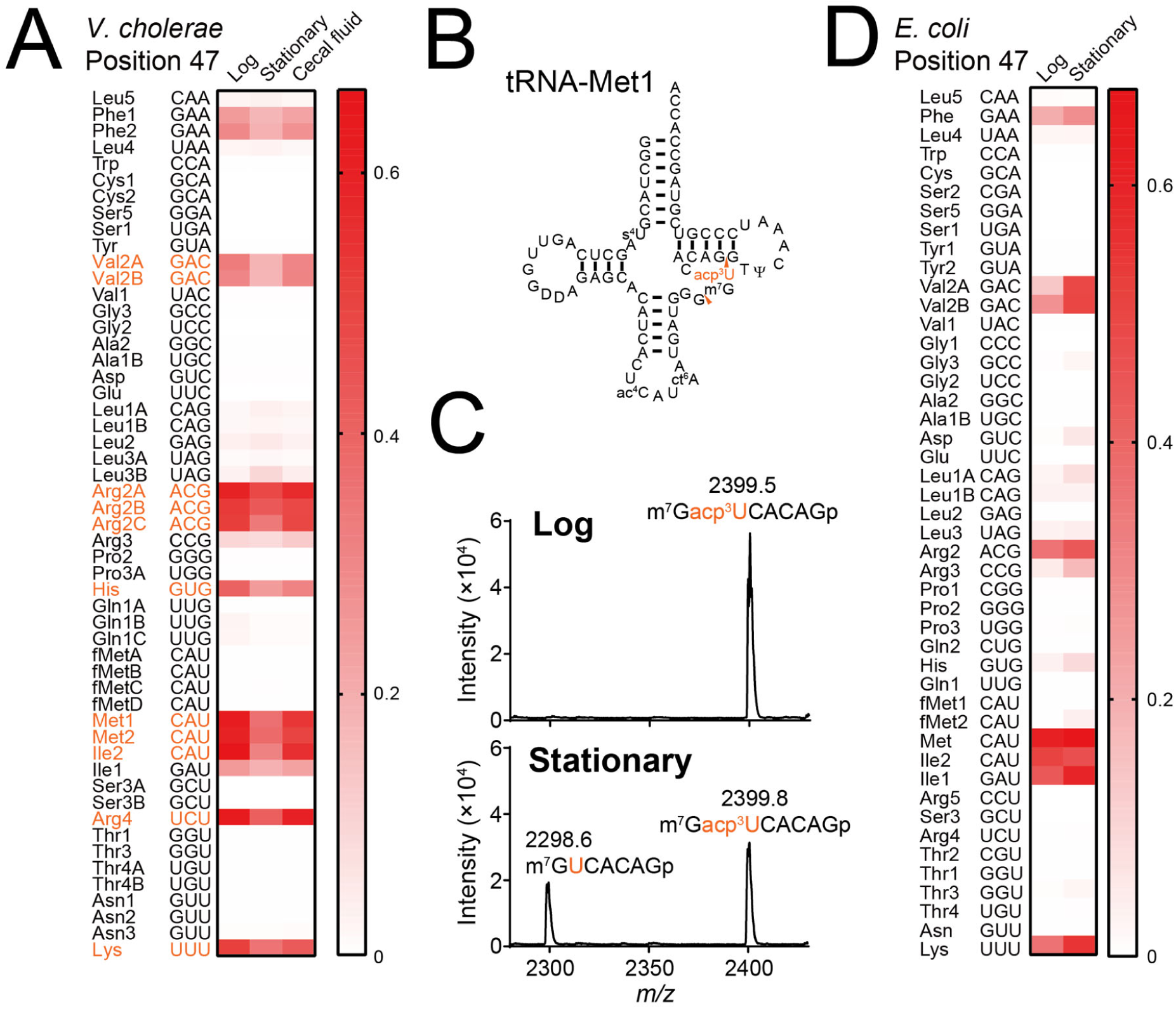
Frequency of tRNA modification by acp^3^U is dependent upon growth phase. (A) Heatmap of misincorporation frequency at position 47 in *V. cholerae* tRNAs isolated from indicated growth condition. Signal intensities in each condition are the average values of three independent tRNA-seq datasets. tRNA species that showed significant differences between signals from log and stationary phase cells are colored in red (multiple two-sided t-test, FDR < 10 %). (B) Secondary structure of tRNA-Met1 with modifications. RNase T_1_ cleavage sites that form the fragment containing position 47 are indicated by red arrowheads. (C) MALDI analysis (positive polarity mode measurements) of RNase A digests of tRNA-Met1 isolated from log and stationary phase samples. Nucleosides at position 47 are colored in red. (D) Heatmap of misincorporation frequency at position 47 in *E. coli* tRNAs isolated from indicated. growth condition. Signal intensities in each condition represent the values of one tRNA-seq dataset.

We hypothesized that RT signatures found in *V. cholerae* but not *E. coli* (Fig. 1AB) could reflect *V. cholerae*-specific modifications, and used RNA mass spectrometric analysis to identify the chemical moieties at some of these positions, focusing initially on U20B in tRNA-Glu and U46 in tRNA-Gln1B. Nucleoside analyses, which reveal the composition of modified ribonucleosides in each tRNA, were carried out on purified tRNA-Glu and tRNA-Gln1B, and revealed that tRNA-Glu contains Ψ, mnm^5^s^2^U, T and Gm, while tRNA-Gln1B contains D, Ψ, cmnm^5^s^2^U, T, s^4^U and m^2^A (Supplementary Fig. 3A and Supplementary Data 3). In addition, these analyses showed that tRNA-Glu and tRNA-Gln1B both also possesses a ribonucleoside whose molecular weight is 387 (Fig. 3A, Supplementary Fig. 6). Since no known modified ribonucleoside has a mass of 387^2^, we postulated that this nucleoside (designated N387) is a novel modification that may be incorporated at various sites (e.g., U20B or U46) within tRNAs. To confirm the positions of N387, we performed fragment analyses. N387-containing fragments were detected using MALDI-TOF mass spectrometry of tRNAs digested with RNase A, which cleaves at the 3’ end of C and U, or RNase T_1_, which cleaves at the 3’ end of G (Fig. 3A and Supplementary Fig. 7). RNase A digests revealed fragments of *m/z* 2121.7 in tRNA-Glu and 1431.1 in tRNA-Gln1B, consistent with the location of N387 at U20B and U46 in these tRNAs, respectively (Fig. 3A). These results suggest that the *V. cholerae*-specific misincorporation signals in tRNA-Glu and tRNA-Gln1B result from N387.

**Figure 3.**
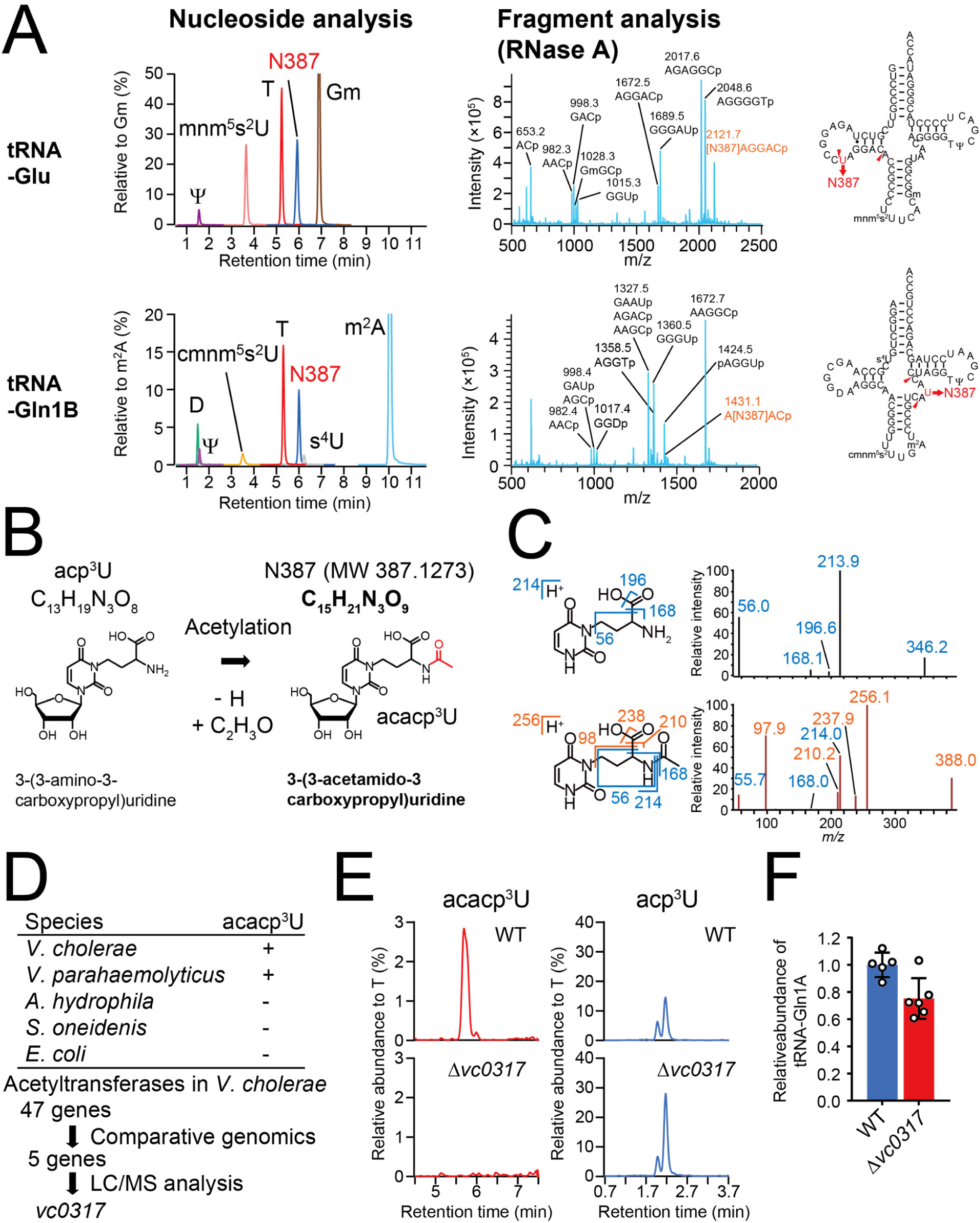
Structure and biosynthesis of acacp^3^U. (A) RNA mass spectrometric analyses of *V. cholerae* tRNA-Glu (Upper) and tRNA-Gln1B (Lower). Left panels show nucleoside analyses by multiple reaction monitoring (MRM), showing the presence of a nucleoside whose mass is 387 (N387), found in neutral loss scans (Supplementary Fig. 6), along with known modifications (denoted in black). The middle panels show fragment analyses of RNase A digests. The fragments containing N387 are colored in red. The right panels show the secondary structures containing modifications based on nucleoside and fragment analyses (Supplementary Fig. 7). (B) Schematic of potential derivation of acacp^3^U (N387) from acp^3^U. (C) MS/MS analyses of acp^3^U (upper panels) and N387 (lower panels). Left panels show the structures and fragmentation patterns of the acp^3^U and acacp^3^U base components. Right panels show the product ion spectra of acp^3^U in tRNA-Met1 (precursor ion; *m/z* 346) and N387 in tRNA-Glu (precursor ion; *m/z* 388). Fragment ions observed in acp^3^U are colored in blue and N387 specific fragment ions are colored in red. (D) Comparative genomic approach to identify an acetyltransferase required for acacp^3^U synthesis. (E) *vc0317* is required for the acetylation of acacp^3^U. Nucleoside analyses detecting acacp^3^U (left) and acp^3^U (right) in WT (upper) and Δ*vc0317* (lower) strains. (F) Abundance of tRNA-Gln1A in WT and Δ*vc0317* strains. tRNAs were quantified through northern blotting and normalized with the abundance of 5S rRNA; and average values, SD, and individual biological replicates (WT; n = 5 and Δ*vc0317*; n = 6) are shown as bars, error bars and circles, respectively (p = 0.01, two-sided t-test).

High-resolution mass spectrometric analysis of N387 from tRNA-Glu yielded a mass value of 387.1273; the best matched chemical formula^10^ corresponding to this mass value is C15H21N3O9 (Fig. 3B). This formula is close to that of acp^3^U (C13H19N3O8), with the difference in chemical composition between these compounds corresponding to acetylation (C2H2O). Since acp^3^U contains a primary amine, a plausible target of acetylation, we predicted that N387 is acetylated acp^3^U, i.e., 3-(3-<u>ace</u>t<u>a</u>mide<u>c</u>arboxy<u>p</u>ropyl)uridine (acacp^3^U). MS/MS analysis with acp^3^U and N387 was used to test this hypothesis. The product ions of acp^3^U, e.g., *m/z* 56, 168 and 214, were also observed in the spectrum of N387, consistent with the presence of acp^3^U in the structure of N387 (Fig. 3C). Several additional fragment ions (e.g., *m/z* 238 and 210) further corroborate the proposed structure of N387 as acacp^3^U. MS/MS spectra of N387 were nearly identical in tRNA-Gln1A and tRNA-Gln1B (Supplementary Fig. 8), suggesting that these tRNAs also contain acacp^3^U.

A comparative genomics approach was then used to identify an acetyltransferase that is required for acacp^3^U formation. We analyzed total tRNA from bacterial species that are phylogenetically close to *V. cholerae* (*Vibrio parahaemolyticus, Aeromonas hydrophila*, and *Shewanella oneidenis*), and found that *V. parahaemolyticus* contains acacp^3^U but the other bacteria do not (Fig. 3D and Supplementary Fig. 9). Based on these results and our *E. coli* data, we identified candidate acetyltransferases in *V. cholerae* (Fig. 3D and Supplementary Data 4). Of the 47 genes annotated as acetyltransferases in the COG database^11^, only 5 are present in *V. cholerae* and *V. parahaemolyticus* but absent in *A. hydrophila, S. oneidenis* and *E. coli* (Fig. 3D and Supplementary Data 4). Nucleoside analysis of total tRNA from transposon insertion mutants corresponding to each of these 5 loci^12^ detected a decreased acacp^3^U signal and an increased acp^3^U signal in tRNA from *vc0317*::Tn (Supplementary Fig. 10). Furthermore, analysis of tRNA from an in-frame *vc0317* deletion mutant revealed that disruption of *vc0317* abolished the acacp^3^U signal and increased the acp^3^U signal (Fig. 3E). Collectively, these results strongly suggest that *vc0317* mediates acetylation during acacp^3^U synthesis and we renamed *vc0317* as *acpA* (for <u>acp</u>^3^U <u>a</u>cetylation). Notably in PFAM^13^, *acpA* is predicted to encode an *N-*acetyltransferase, providing further support for the idea that N387 includes acetylation of a primary amine group. One effect of acetylation of acp^3^U involves tRNA abundance, with modestly reduced tRNA-Gln1A levels in log phase cultures of Δ*acpA* compared to the wild type strain (Fig. 3F).

*V. cholerae* tRNA-seq analysis also revealed a high misincorporation frequency at C32 in tRNA-Tyr (Fig. 1A). Most sequencing reads contained U rather than C at position 32, even though C is present in all 5 copies of this tRNA gene^14^ (Supplementary Fig. 11). RT-PCR coupled with direct Sanger sequencing confirmed the presence of this C-to-U conversion in tRNA-Tyr (Fig. 4A). However, ribonucleoside and fragment analysis of purified tRNA-Tyr did not detect any modifications that could be assigned to position 32, although it successfully assigned several other modifications (Fig. 4B, Supplementary Fig. 12). Instead, fragment analysis revealed a U32-containing fragment (AGAUp, *m/z* 1326.25) (Fig. 4C), strongly suggesting that C32 undergoes post-transcriptional C-to-U RNA editing. Intriguingly, tRNA-Tyr from a strain lacking *miaA*, which is responsible for the initial step of biosynthesis of ms^2^io^6^A at position 37 in tRNA-Tyr, retains C at position 32 (Fig. 4AC). tRNA-Tyr from a strain lacking *thiI*, whose product synthesizes s^4^U at position 8 and 9 in tRNA-Tyr, contained both C and U at position 32 (Fig. 4A). These results suggest that C-to-U editing at this site depends upon the presence of other modification(s) or the associated modification enzymes. The enzyme responsible for C-to-U RNA editing in *V. cholerae* remains unknown. Since *T. brucei’s* A-to-I editing enzyme is reported to catalyze C-to-U editing as well^15^, it is possible that VC0864, *V. cholerae*’s presumed A-to-I tRNA editing enzyme, could also be involved in C-to-U editing

**Figure 4.**
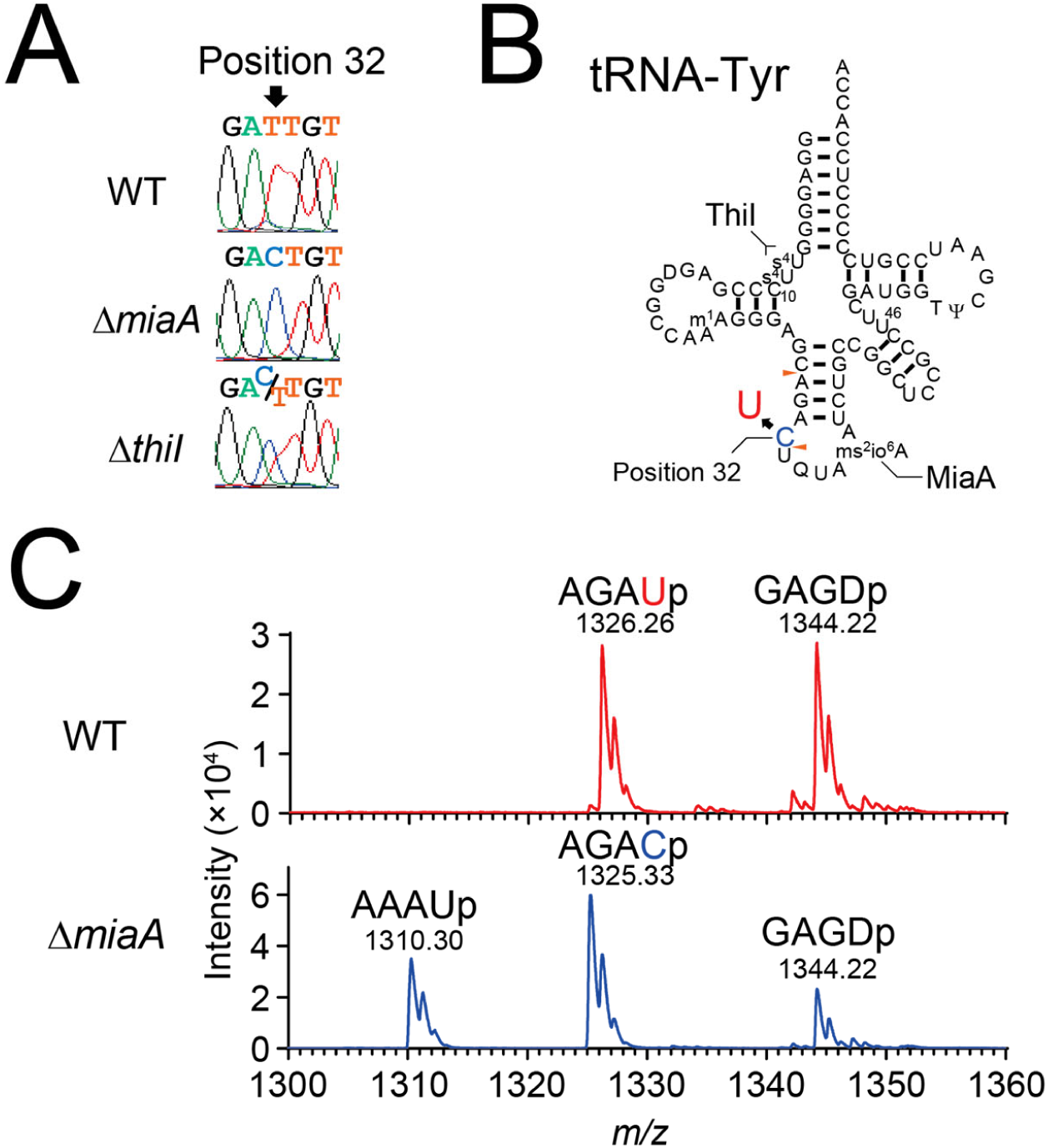
Cytidine at position 32 in tRNA-Tyr undergoes C-to-U RNA editing. (A) Sanger sequencing of cDNA of tRNA-Tyr from WT (top), Δ*miaA* (middle), and Δ*thiI* (bottom) strains. Position 32 is indicated by the arrow. (B) Secondary structure of tRNA-Tyr with modifications based on fragment analyses (Supplementary Fig. 12). s^4^U and i^6^A, which is a precursor of ms^2^io^6^A, are synthesized by *thiI* and *miaA*, respectively. RNase A cleavage sites that form the fragment containing position 32 are indicated by red arrowheads. (C) Fragment analyses of an oligo protected portion (position 10 to 46) of tRNA-Tyr from WT (upper) and Δ*miaA* (lower) strains. *m/z* values of detected peaks with assigned fragment sequences are shown. The MALDI analyses were conducted in negative polarity mode.

Our findings demonstrate that comparative tRNA-seq provides a high-throughput method for cataloging sites of likely tRNA modification and identifying tRNA species or conditions warranting more in-depth analyses. By combining tRNA-seq and RNA mass spectrometry (tRNA-SMS) to characterize *V. cholerae’s* tRNA, we uncovered a species-specific modification (m^1^A) and discovered a new tRNA modification (acacp^3^U) along with an enzyme required for its synthesis. Moreover, this approach yielded evidence of C-to-U RNA editing, a process not previously observed in bacteria. Thus, our data reveals substantial diversity in tRNA modifications among even phylogenetically closely related organisms like *V. cholerae* and *E. coli*. Since the full complement of tRNA modifications have been well characterized in only a few model organisms, e.g., *E. coli* and *Saccharomyces cerevisiae*, it is likely that there is a plethora of as yet undescribed modifications. The tRNA-SMS approach should be useful to probe the diversity of tRNA modifications throughout all three kingdoms of life.

## Methods

### Strains and culture conditions

The strains used in this study are listed in Supplementary Table 2. *V. cholerae* C6706, a clinical isolate^16^, and *E. coli* MG1655 were used in this study as wild-type strains. All *V. cholerae, E. coli*, and *V. parahaemolyticus* strains were grown in LB containing 1 % NaCl at 37 °C. *E. coli* SM10 (lambda pir) harboring derivatives of pCVD442^17^ was cultured in LB plus carbenicillin (Cb). Antibiotics were used at the following concentrations: 200 μg/mL streptomycin, 50 μg/mL Cb. *Aeromonas hydrophila* and *Shewanella oneidensis* were cultured at 30°C in nutrient broth (BD) and Tryptic Soy Broth, respectively.

### Strain construction

All mutations in C6706 were created using homologous recombination and a derivative of the suicide vector pCVD442^17^. Targeting vectors for gene deletions contained ∼1000 bp of DNA flanking each side of the target gene cloned into pCVD442’s SmaI site using isothermal assembly.

### RNA extraction

Total RNA was extracted with TRIzol (Life Technologies) according to the manufacturer’s instructions. The tRNA fraction was cut out from 10 % TBE-Urea gels and recovered by isopropanol precipitation.

### Isolation of individual tRNA species

One liter cultures of log-phase (OD_600_ = 0.3) and stationary phase (24h) *V. cholerae* cells were harvested, and total RNA was extracted^18^. Briefly, cells were resuspended in 5 mL buffer [50 mM NaOAc, pH 5.2, 10 mM Mg(OAc)_2_], mixed with 5 mL water saturated phenol, and agitated vigorously for 1 h. The aqueous phase was separated by centrifugation, washed with chloroform, and recovered by isopropanol precipitation. RNA was run through a manually packed DEAE column (GE healthcare) to remove contaminants and recovered by isopropanol precipitation. Individual tRNA species were bound to biotinylated DNA probes anchored to high-capacity streptavidin agarose resin (GE Healthcare) in 30 mM Hepes-KOG, pH 7.0, 1.2 M NaCl, 15 mM EDTA, and 1 mM DTT at 68 °C for 30 min with shaking. Beads were washed three times with 15 mM Hepes-KOH, pH 7.0, 0.6 M NaCl, 7.5 mM EDTA, and 1 mM DTT and seven times with 0.5 mM Hepes-KOH, pH 7.0, 20 mM NaCl, 0.25 mM EDTA, and 1 mM DTT. Purified tRNAs were extracted from beads with TRIzol. After Turbo DNase (Thermo Fisher Scientific) treatment to remove residual DNA probes, purified tRNAs were purified on 10 % TBE-Urea gels. The probes used in this study are listed in Supplementary Data 5.

### tRNA sequencing

Total tRNA fraction (250 ng) was deacylated in 500 μl of 100 mM Tris-HCl pH 9.0 at 37 °C for 1 hr and recovered by isopropanol precipitation. After dephosphorylation with alkaline phosphatase from calf intestine (New England Biolabs), tRNAs were ligated to 100 pmol of 5’ adenylated and 3’ end-blocked DNA oligo (3’ linker, Supplementary Data 5) using truncated T4 RNA ligase at 25 °C for 2.5 hr in 25 % PEG 8000. The ligated product was purified on a 10 % TBE-Urea polyacrylamide gel (Thermo Fisher Scientific) as above. Half of the recovered ligated tRNAs were reverse transcribed with 5 pmol TGIRT-III (InGex) in 100 mM Tris-HCl pH 7.5, 0.5 mM EDTA, 450 mM NaCl, 5 mM MgCl_2_, 5 mM DTT, 1 mM dNTPs, and 1.25 pmol primer (ocj485, Supplementary Data 5) at 60 °C for 1 hr. After the elimination of template RNAs by alkali treatment, cDNA was purified on a 10 % TBE-Urea polyacrylamide gel. The single stranded cDNA was then circularized using 50 U of CircLigase II(Epicenter) at 60 °C for 1 hr, followed by addition of another 50 U of CircLigase II for an additional 1 hr at 60 °C. cDNA was amplified using Phusion DNA polymerase (New England Biolabs) with o231 primer and index primers (Supplementary Table S2). After 12-18 rounds of PCR amplification, the product was gel purified from an 8 % TBE-Urea polyacrylamide gel (Thermo Fisher Scientific). Sequencing was performed using a Illumina miSeq. 3’ linker sequences and one nucleotide at the 5’ end was trimmed. Bowtie^19^ v. 1.2.2 with default settings was used for mapping reads to reference tRNA sequences (Supplementary Data 6) retrieved from tRNAdb^7^. Two sequences in *V. cholerae* (tdbD00003706, tdbD00008082) and three sequences in *E. coli* (tdbD00007320, tdbD00010329, and tdbD00011810) were eliminated from the reference sequences due to extremely low coverage. Mpileup files were made using the samtools mpileup command without any filtration (option, -A –ff 4 -x -B -q 0 - d 10000000 -f). The frequency of misincorporation was calculated in each mpileup file. 5’ end termini of the mapped reads were piled up using the bedtools genomecov command (-d -5 -ibam). To calculate the termination frequency, the number of 5’ termini at any given position was divided by the total number of mapped termini at the given position along with all upstream positions (5’ side). Frequencies of misincorporation and termination of one replicate were visualized with R (3.4.3) or Graphpad Prism.

Reference sequences of *E. coli* tRNAs along with the catalogue of modifications in the tRNA database and the literature^7,20-27^ (Supplementary Data 1 and Supplementary Table 1) were used to assign modifications to the *E. coli* tRNA-seq data in Supplementary Figure 2. Misincorporation frequencies of > 5% at sites of modification in the reference database were used as a threshold for assignment of predicted modifications. Termination frequencies of >5% were also used as a threshold for assignments of modified sites; however, since termination signals were usually detected two nucleotides downstream from known modified sites, assignments were adjusted accordingly (except for DD, acp^3^U, k^2^C, and m^6^t^6^A (see Supplementary Table 1)). E. coli modifications were considered to be predictable when signals (either of misincorporation or termination) were present in ≥ 50% of known modified sites.

*V. cholerae* modifications were also assigned with >5% thresholds for misincorporation or termination. Although G at position 10 (G10) in several tRNAs and G at position 35 (G35) in tRNA-Leu2 have misincorporation signals greater than 5%, they were excluded from further analysis because these signals were also observed in the *E. coli* data, where G10 and G35 modifications have not been reported. These elevated misincorporation signals may arise from other factors including residual secondary or tertiary structures or specific sequence contexts influencing reverse transcription.

### Northern blotting

In total, 0.3 μg RNA was electrophoresed on 10 % Novex TBE-Urea gels (Thermofisher) and stained with SYBR Gold (Life Technologies). RNA was transferred to nitrocellulose membranes by semidry blotting and cross-linked twice to membranes with 1200 μJ UV light. Membranes were incubated in ULTRAhyb-oligo (Life Technologies) at 42 °C for 30 min followed by hybridization overnight at 42 °C with 4 pmol DNA probes radiolabeled with [γ-^32^P]ATP (PerkinElmer) and T4 Polynucleotide kinase (New England Biolabs). Membranes were washed twice with 2 × SSC/0.5 % SDS, and then bound probe was detected using an FLA-5000 phosphoimager (Fuji). The signal intensity of tRNA was normalized to that of 5S rRNA. All DNA oligos were synthesized by Integrated DNA Technology. Probe sequences are listed in Supplementary Data 5.

### Nucleoside analysis

100 ng of total RNAs or isolated tRNAs were digested with 0.5 unit Nuclease P1 and 0.1 unit of phosphodiesterase I in 22 µl reactions containing 50 mM Tris-HCl pH 5.3, 10 mM ZnCl_2_ at 37 °C for 1 h. Reaction mixtures were then mixed with 2 μl 1M Tris-HCl pH 8.3 and 1 μl of 1 unit/μl Calf Intestine phosphatase and incubated at 37 °C for 30 min. Enzymes were removed by filtration using 10 K ultrafiltration columns (VWR). 18 μl aliquots were mixed with 2 μl of 50 μM ^15^N-dA and 2.5-10 μl of digests were injected into a Agilent 1290 uHPLC system bearing a Synergi Fusion-RP column (100 × 2 mm, 2.5 μm, Phenomenex) at 35 °C with a flow rate 0.35 ml/min with a solvent system consisting of 5 mM NH_4_OAc (Buffer A) and 100 % Acetonitrile (Buffer B). The gradient of acetonitrile was as follows: 0 %; 0-1 min, 0-10 %; 1-10 min, 10-40 %; 10-14 min, 40-80 %; 14-15 min, 80-100 %; 15-15.1 min, 100 %; 15.1-18 min, 100-0 %; 18-20 min, 0 %; 20-26 min. The eluent was ionized by an ESI source and directly injected into a Agilent 6460 QQQ. The voltages and source gas parameters were as follows: gas temperature; 250 °C, gas flow; 11 L/min, nebulizer; 20 psi, sheath gas temperature; 300 °C, sheath gas flow; 12 L/min, capillary voltage; 1800 V, and nozzle voltage; 2000 V.

Dynamic multiple reaction monitoring (MRM) was carried out to survey known modifications. In the first quadrupole, a proton adduct of a target nucleoside was selected as a precursor ion based on its mass to charge ratio (*m/z*). Only singly charged ions, i.e., z equals 1, were observed. In the second quadrupole, the precursor was broken by collision inducible dissociation (CID) to produce nucleoside species specific product ions, which in many cases were proton adducts of different bases. Then, one specific product ion was selected in the third quadrupole based on its *m/z* value and delivered to the detector. This multiple selection approach, with targeting of specific precursor and product ions, enables high signal to noise ratios. The retention time windows and *m/z* values of precursor and product ions for dynamic MRM analyses are listed in Supplementary Data 3.

The neutral loss scan (NLS) method was used to search for unknown modifications. For one run, we used ∼50 mass values of precursor ions with 1 Da intervals. Then, a mass of the product ion was set 132 Da lower than that of the precursor ion, a value corresponding to the loss of a ribose moiety. We performed five runs to cover precursor ions whose *m/z* values ranged from 244 to 445 (Supplementary Fig. 6). The presence of N387 was then confirmed by MRM analysis.

For MS/MS analysis, 1 μg of isolated tRNAs were hydrolyzed. In this analysis, we selected singly-protonated ions with *m/z* 388 for N387 and 346 for acp^3^U as precursor ions in the first quadrupole, respectively; after CID in the second quadrupole, an *m/z* scan from 10 to 1000 was carried out in the third quadrupole, yielding the mass spectra of the fragments.

To measure the mass of N387 precisely, 2.5 μg of tRNA-Glu was digested as described above and 500 ng of the digest was subjected to the HPLC system described above coupled with an Agilent 6520 quadrupole time-of-flight (QTOF) mass spectrometer. An offset value that is an average error ppm value of 5.9 ppm in known nucleosides (including A, G, Gm, mnm^5^s^2^U, and m^2^A), was added to the measured mass value (387.1251) to obtain the calibrated mass value of N387 (387.1274). The chemical formula was explored in ChemCalc Molecular Formula Finder^10^ with the following constraints: C,9-100, H,0-100, N,2-10, O,5-10, S,0-3, unsaturation,3-8, with restriction to integral unsaturation values.

### Fragment analysis

400-1000 ng isolated tRNAs were digested in 3 μl aliquot with 20 ng RNase A (QIAGEN) in 10 mM NH_4_OAc pH 7 or 20 unit RNase T_1_ in 10 mM NH_4_OAc pH 5.3 at 37 °C for 1 hr. On a MALDI steel plate, 0.5 μl of matrix (0.7 M 3-hydroxypicolinic acid (HPA) and 70 mM ammonium citrate in 50 % milliQ and 50 % acetonitrile) was mounted and dried, followed by mounting of 0.5 μl RNase digests and drying. The samples were analyzed with Bruker Ultraflex Xtreme MALDI-TOF mass spectrometer.

### Oligo protection

2.8 and 3.8 μg of tRNA-Tyr from WT and Δ*miaA* were mixed with 500 pmol of DNA oligos (Supplementary Data 5) in 50 μl aliquots containing 50 mM Hepes KOH pH 7.6, 150 mM KCl and heated to 90 °C for 1 min and gradually cooled down to room temperature at 1 °C/min for annealing, followed by RNase digestion with 50 ng RNase A and 50 unit RNase T_1_ on ice for 15 min. Protected DNA/RNA duplexs were purified on 10 % TBE-Urea gels and recovered by isopropanol precipitation and dissolved in 5 μl milliQ. A 2 μl aliquot was subjected to fragment analysis with RNase A using a MALDI-TOF spectrometer as described above.

### Comparative genomics

Protein sequences were retrieved from NCBI (*V. cholerae*; GCF_000006745.1. *V. parahaemolyticus*; GCF_000196095, *E. coli*; GCF_000005845.2, *S. oneidenis*; GCF_000146165.2, and *A. hydrophila*; GCF_000014805.1). The *V. cholerae* protein sequences were queried with the other species’ protein sequences with local BLAST and an E-value threshold of 1E-10 to identify similar sequences. Forty seven *V. cholerae* proteins that include “acetyltransferase” in their COG names were retrieved from COG 2003-2014 update using R^11^. Five of these putative acetyltransferases were found to be present in *V. cholerae* and *V. parahaemolyticus*, but not in *A. hydrophila, S. oneidenis* and *E. coli* when 1E-10 was used as a threshold.

### Infant rabbit infection

Mixed-gender litters of 2 day old New Zealand white infant rabbits, cohoused with a lactating mother (Charles River) were inoculated with wild-type *V. cholerae* as described^9^. Approximately 20hr post-inoculation, the rabbits were sacrificed and *V. cholerae* in the cecal fluid was collected by centrifugation. Total RNA was then extracted using TRIzol reagent.

### Animal use statement

Infant rabbit studies were conducted according to protocols approved by the Brigham and Women’s Hospital Committee on Animals (Institutional Animal Care and Use Committee protocol number 2016N000334 and Animal Welfare Assurance of Compliance number A4752-01) and in accordance with the recommendations in the Guide for the Care and Use of Laboratory Animals of the Nathional Institutes of Health and the Animal Welfare Act of the U.S. Department of Agriculture.

## Data availability

All data is available from the corresponding authors upon request.

## Code availability

All codes are available from the corresponding authors upon request.

## Competing interests

The authors declare no competing interests.

## Acknowledgments

We thank Brigid Davis, Troy Hubbard and Waldor lab members for helpful comments on the project and the manuscript, and the Harvard FAS Science Core Facility for use of MALDI equipment. This work was supported by NIH R01-AI-042347 and HHMI (MKW)

