## Supplementary Information for "Surveying the landscape of tRNA modifications by combining tRNA sequencing and RNA mass spectrometry"

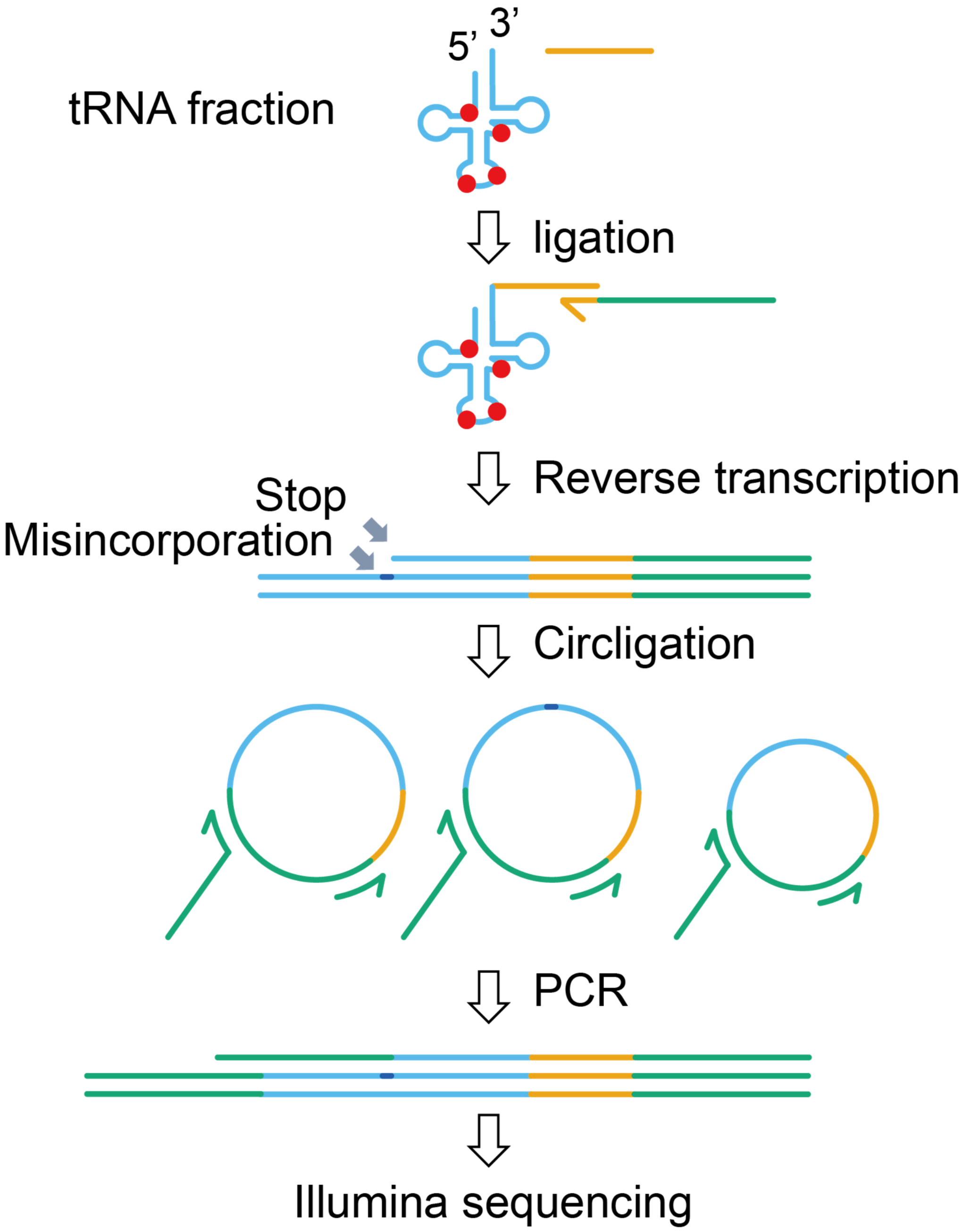


**Supplementary Fig. 1. Scheme of tRNA-seq protocol.**

tRNA was isolated via gel purification from total RNA samples. Following tRNA deacylation, linker DNA (yellow) was ligated to the 3' end of tRNAs. Then, the TGIRT enzyme was used for reverse transcription (green and yellow primer). In this process, tRNA modifications (red circles) may be recorded in cDNAs as an interruption (stop) in reverse transcription and/or by incorporation of mismatched bases (misincorporation). cDNAs are then circularized using Circligase II, followed by PCR. Amplified cDNAs are then subjected to Illumina sequencing.

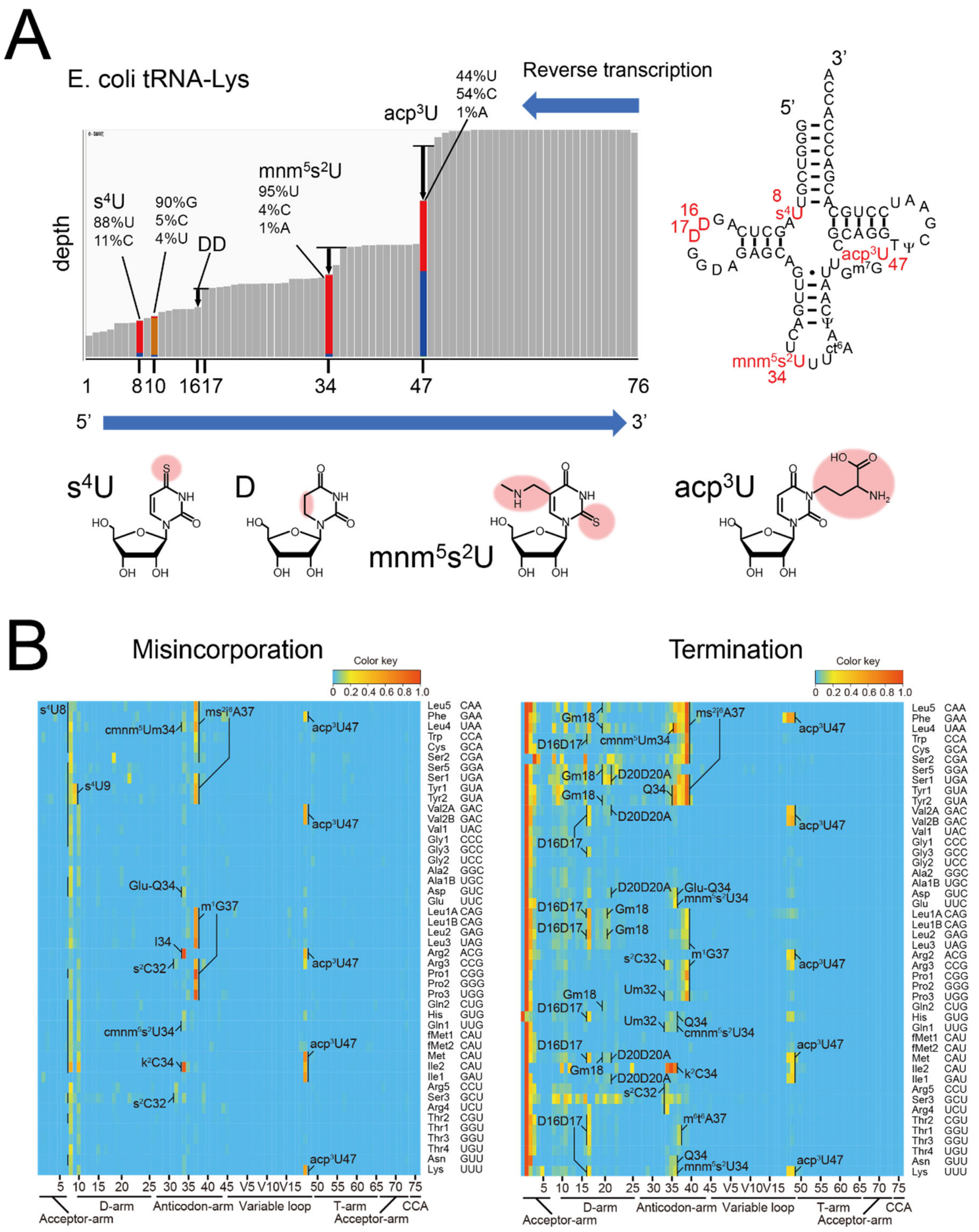


**Supplementary Fig. 2. tRNA-seq profiling of tRNA modifications in *E. coli*.**

(A) Example of the analysis of reverse transcription derived signatures in tRNA-seq data. The bars in the left panel represent mapped read depth with the left side and right side corresponding to the 5' and 3' end of tRNAs, respectively. Bars in which the misincorporation frequency is less than 1% are colored in grey; additional colors are shown at sites where there are higher levels of misincorporation (red corresponds to U, blue, C, orange, G and yellow, A). The significance of the misincorporation signal at position 10 is not known. Several drops in depth around sites of known modifications are also apparent

(e.g. DD at positions 16, 17); however, the correspondence between the decrease in read depth and sites of modification is less precise than with nucleotide misincorporation. The right panel shows the secondary structure of tRNA-Lys. The structures of some of the modifications that lead to reverse transcription derived signatures are shown in lower panels.
(B) Heatmaps of the frequency of misincorporation (left) and stop (termination) of reverse transcription (right) signals in tRNA from stationary phase *E. coli*. Each row represents an individual tRNA and each column represents a position within tRNAs. The modifications are assigned based on the reference tRNA sequences (Supplementary Data 1 and Supplementary Table 1)

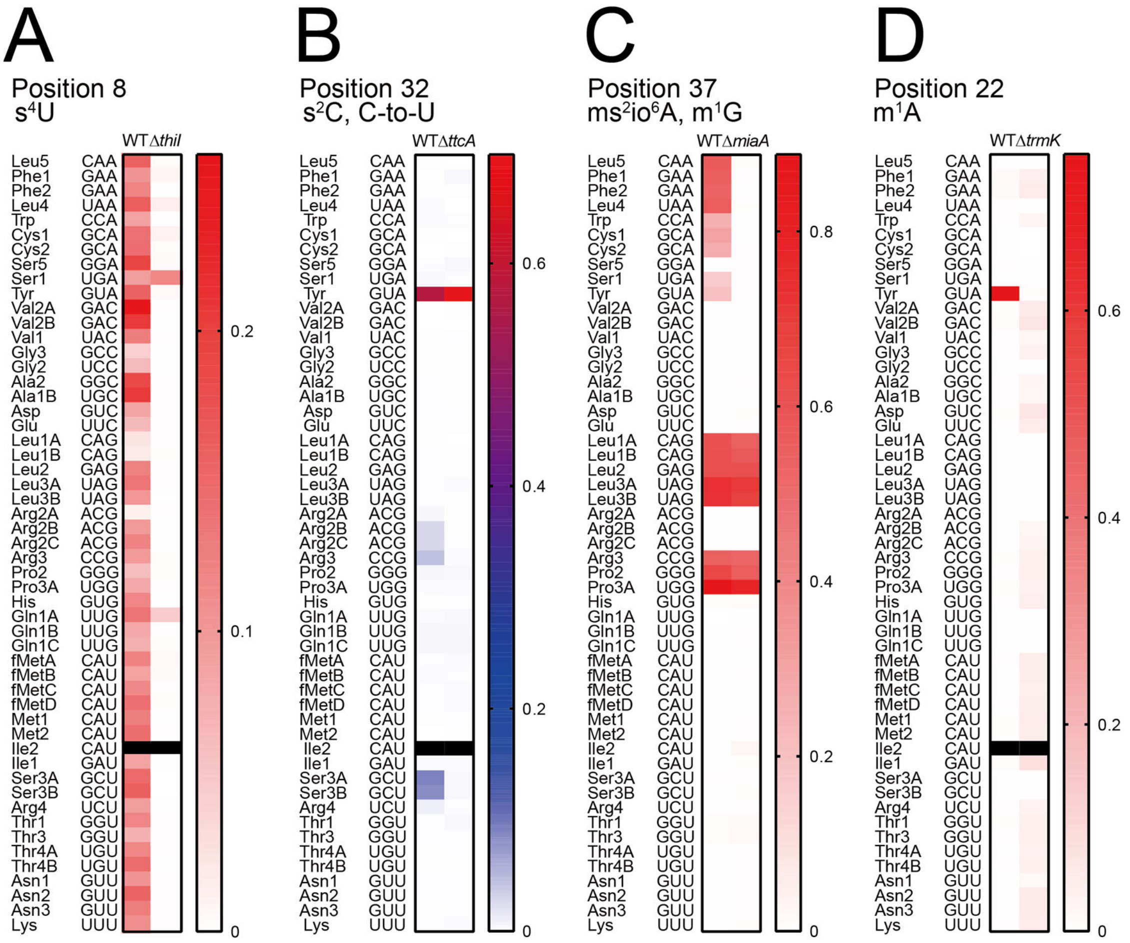


**Supplementary Fig. 3. Validation of modifications inferred from RT-derived signals using mutant *V. cholerae* strains.**

(A) Heatmap of misincorporation frequency at position 8 in *V. cholerae* tRNAs. Most of the misincorporation signals, except for tRNA-Ser1 and tRNA-Gln1A, are eliminated in the Δ*thiI* strain, consistent with the idea that misincorporation results from the associated modification, s$^4$U. The data for tRNA-Ile2 is not shown (black) because of insufficient read depth (<100 reads).

(B) Heatmap of misincorporation frequency at position 32 in *V. cholerae* tRNAs. The signals in tRNAs that are expected to have s$^2$C (tRNA-Arg2A, tRNA-Arg2C, tRNA-Arg3, tRNA-Ser3A, tRNA-Ser3B, and tRNA-Arg4) are eliminated in the Δ*ttcA* strain, whereas the signal in tRNA-Tyr remains due to C to U RNA editing (see Fig 4). The data for tRNA-Ile2 is not shown (black) because of insufficient read depth.

(C) Heatmap of misincorporation frequency at position 37 in *V. cholerae* tRNAs. The signals in tRNAs that are expected to have ms$^2$io$^6$A (tRNA-Leu5, tRNA-Phe1, tRNA-Phe2, tRNA-Leu4, tRNA-Trp, tRNA-Cys1, tRNA-Cys2, tRNA-Ser1, and tRNA-Tyr) are eliminated in the Δ*miaA* strain, whereas the signals in tRNA species that are predicted to have m$^1$G at position 37 remain.

(D) Heatmap of misincorporation frequency at position 22 in *V. cholerae* tRNAs. The signal in tRNA-Tyr is absent in the Δ*trmK* strain, suggesting that this signal is derived from m$^1$A. The data for tRNA-Ile2 is not shown (black) because of insufficient read depth.

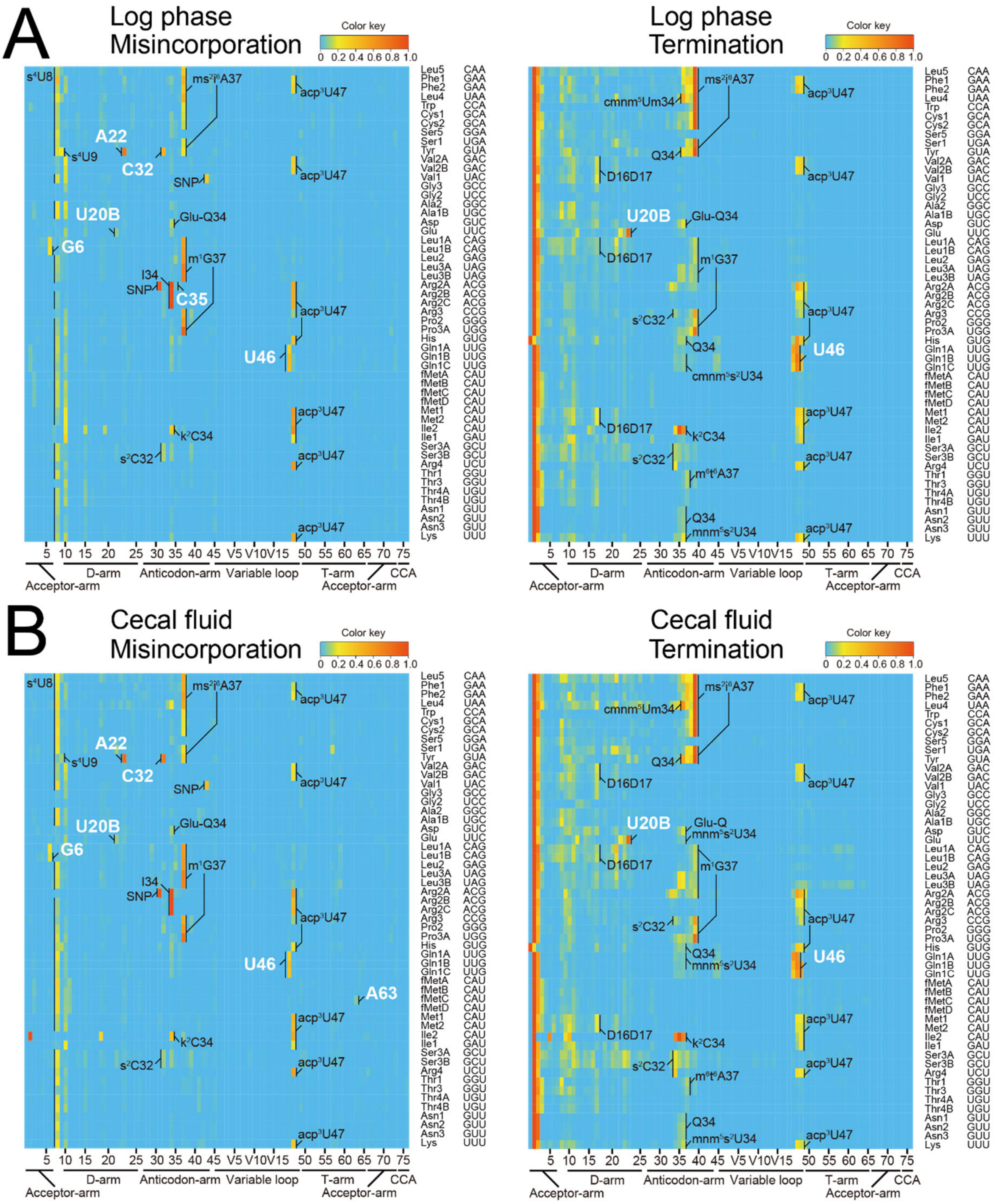

**Supplementary Fig. 4. tRNA-seq profiles of log phase (A) and cecal fluid-derived (B)** ***V. cholerae*.** Heatmaps of frequency of misincorporation (left) and termination of reverse transcription (right) signals in indicated samples. Types and positions of modifications that are presumed shared with *E. coli* are shown in black. Positions of *V. cholerae*-specific signals are indicated in white letters.

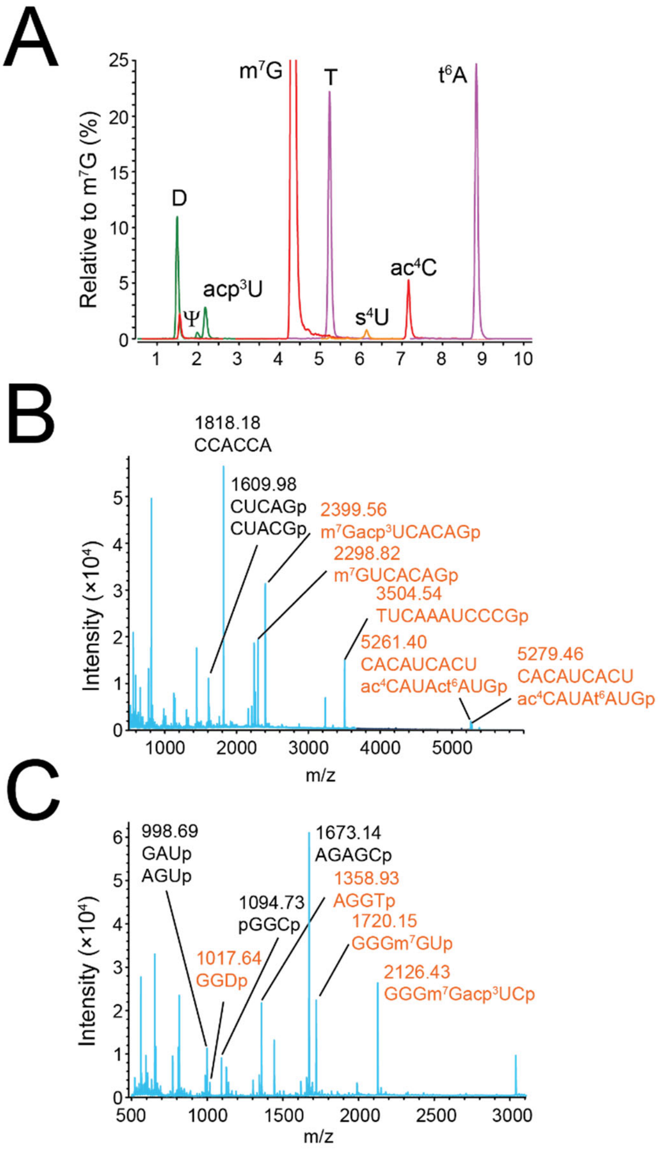


**Supplementary Fig. 5. RNA mass spectrometric analyses of *V. cholerae* tRNA-Met1 from stationary phase.**

(A) Nucleoside analysis detecting D, Ψ, acp$^3$U, m$^7$G, T, s$^4$U, ac$^4$C and t$^6$A.

(B) Fragment analysis of RNase $T_1$ digests. The fragments with or without modifications are shown in red and black, respectively. Measurement was conducted in the positive polarity mode.

(C) Fragment analysis of RNase A digests. The fragments with or without modifications are shown in red and black, respectively. Measurement was conducted in the positive polarity mode.

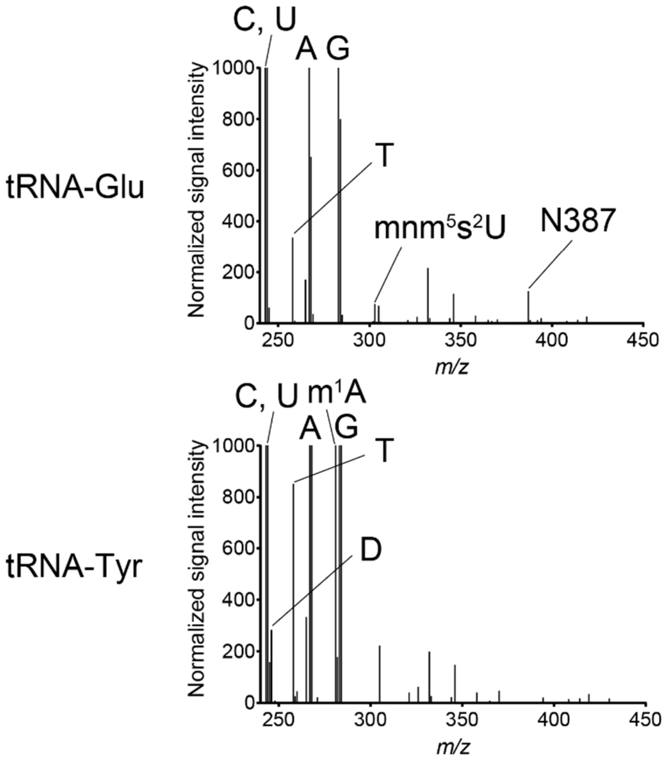


**Supplementary Fig. 6. Neutral loss scan analyses of tRNA-Glu and tRNA-Tyr.**

Nucleosides derived from purified tRNA-Glu and tRNA-Tyr were subjected to LC/MS/MS and product ions that had a mass reduction of 132 Da, corresponding to loss of a ribose moiety, were monitored. The normalized signal intensities (y-axis) were calculated by determination of the total amount of conventional nucleosides and a signal derived from $^{15}$N-dA, which was spiked into each sample. Conventional nucleosides and modified nucleosides, including N387, were assigned by mass values.

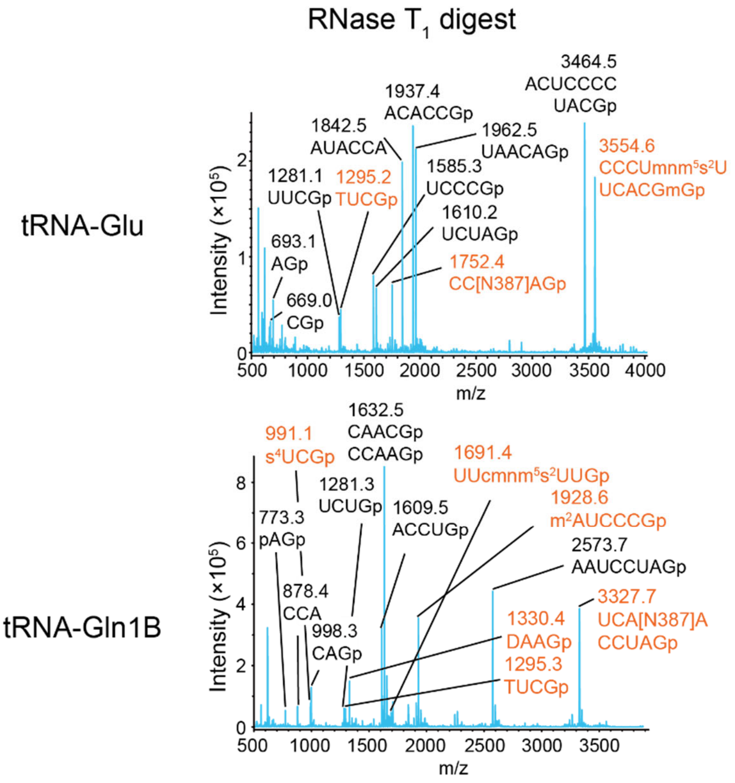


**Supplementary Fig. 7. Fragment analyses of RNase $T_1$ digests of tRNA-Glu (upper) and tRNA-Gln1B (lower).**

The fragments with or without modifications are shown in red and black, respectively. Measurement was conducted in the positive polarity mode.

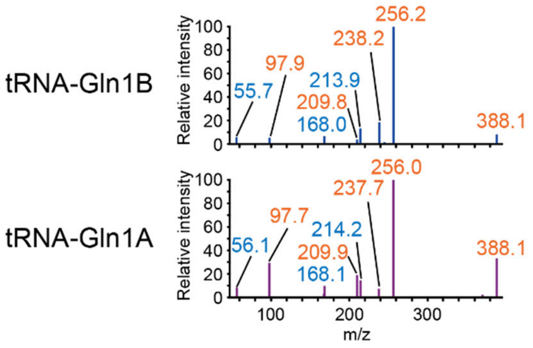


**Supplementary Fig. 8. MS/MS analyses of N387 in tRNA-Gln1B (upper) and tRNA-Gln1A (lower).** Fragment ions observed in $acp^3U$ are colored in blue and N387 specific fragment ions are colored in red.

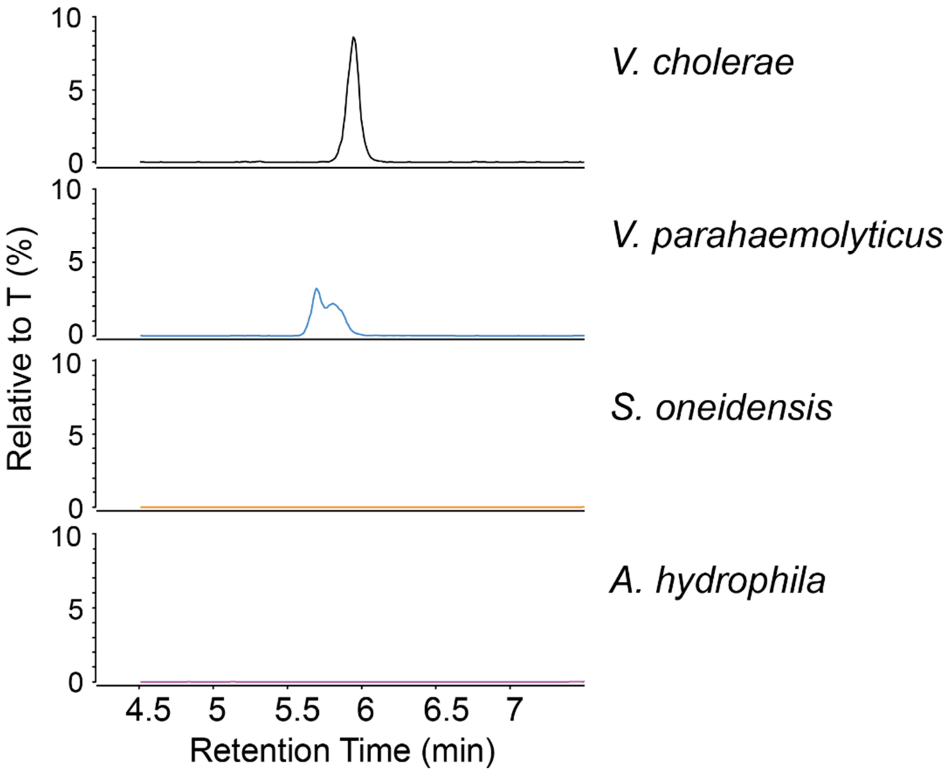


**Supplementary Fig. 9. Presence of acacp$^3$U in *V. cholerae* related organisms.**
Nucleoside analyses of total tRNA fraction from the indicated organisms. Mass chromatograms detecting acacp$^3$U are shown.

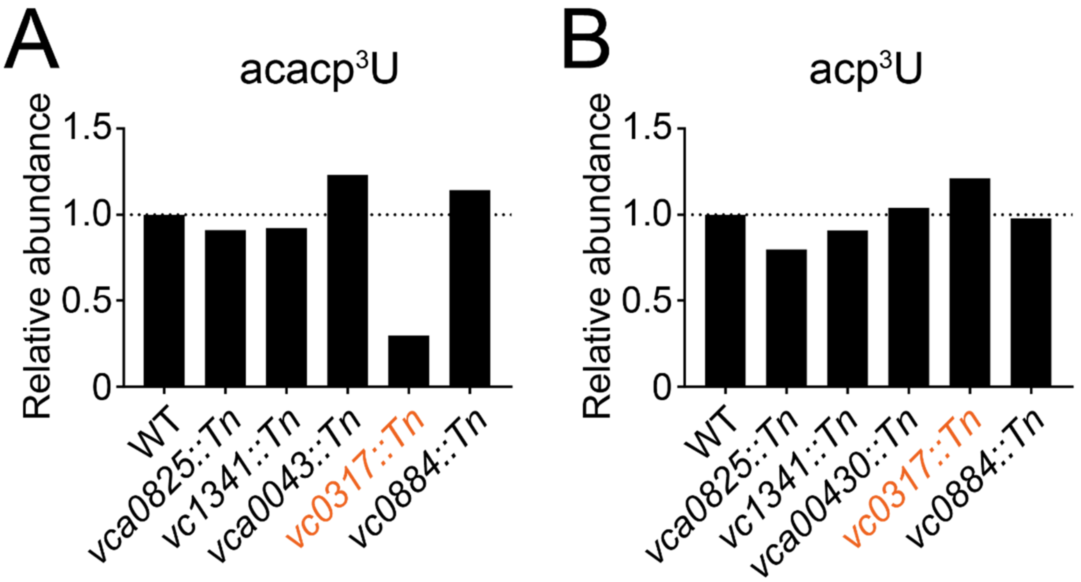


**Supplementary Fig. 10. VC0317 is a candidate acetyltransferase required for acacp³U synthesis.** Nucleoside analysis of total tRNAs derived from strains containing transposon insertions in putative acetyltransferases. Relative abundances of acacp³U (A) and acp³U (B), normalized to that of T, are shown.

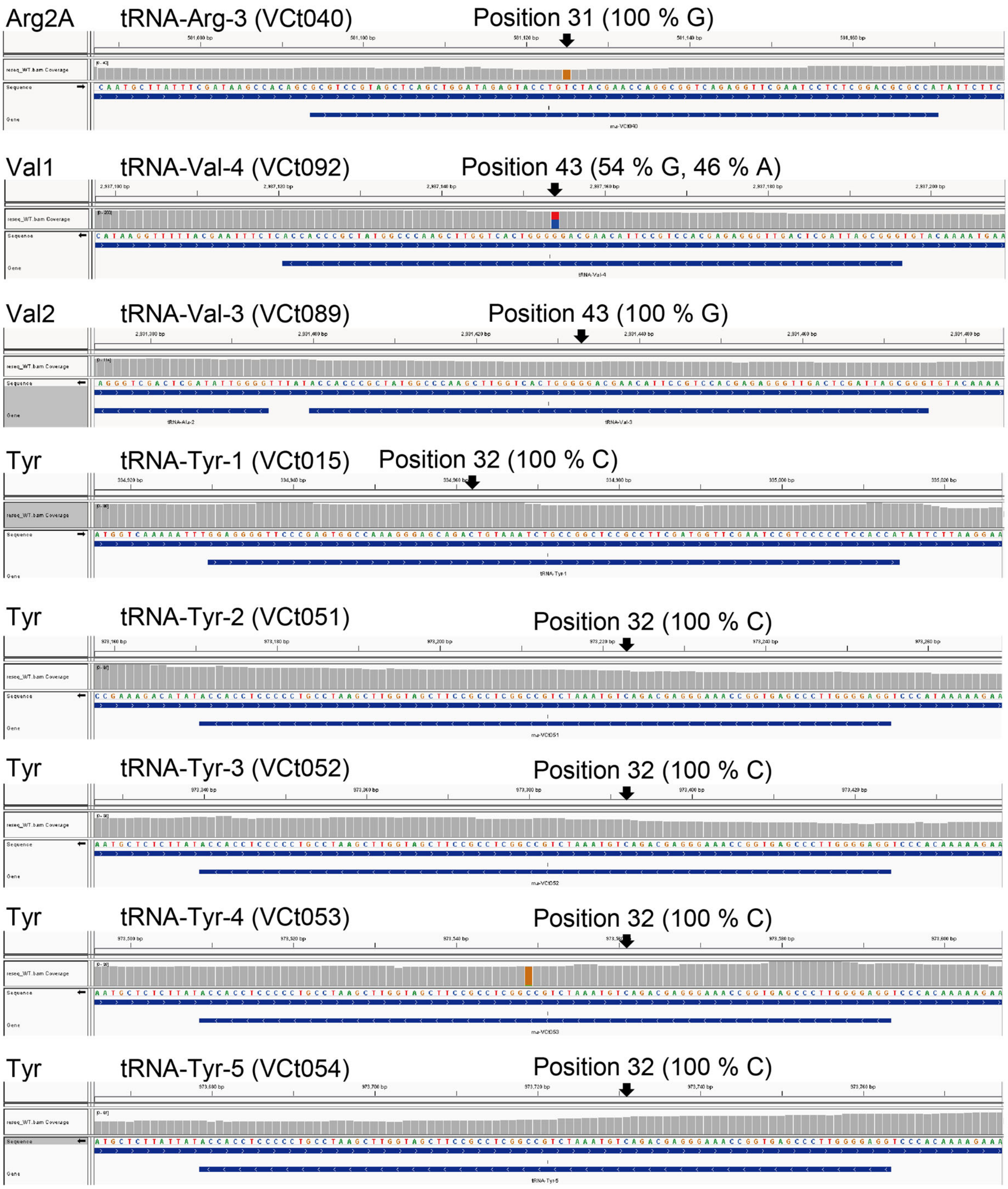


**Supplementary Fig. 11. Genomic sequences of *V. cholerae* tRNAs**

*V. cholerae* genomic tRNA sequences are retrieved from whole genome sequencing data[1]. In contrast to the N16961reference genome[2], in C6706, tRNA-Arg2 has a G (rather than a T) at position 31. Also, one of the two tRNA-Val1 genes has 46% A at position 43 (rather the G), while the other tRNA-Val1 is 100% G at this position, suggesting that a SNP is present in one of the tRNA-Val1 genes. All the tRNA-Tyr genes have 100 % C at position 32.

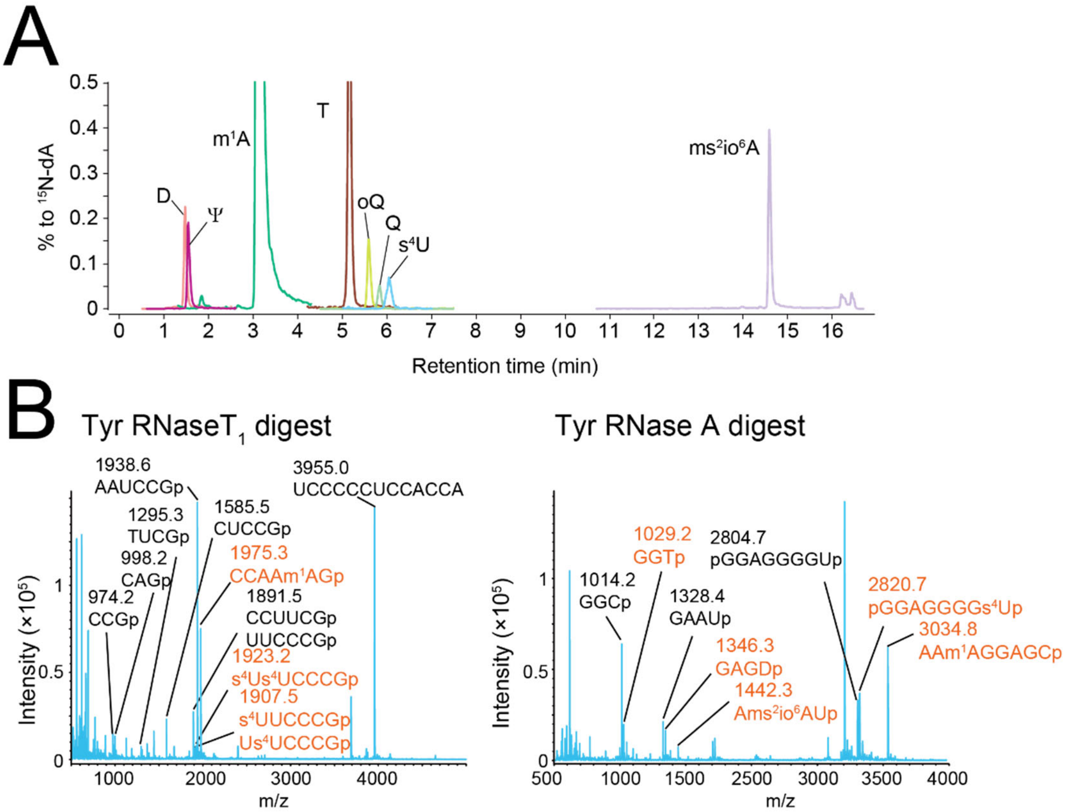


**Supplementary Fig. 12. RNA mass spectrometric analyses of tRNA-Tyr.**

(A) Nucleoside analysis detecting D, Ψ, $m^1A$, T, oQ, Q, $s^4U$, $ms^2io^6A$.

(B) Fragment analyses of RNase $T_1$ (left) and RNase A (right) digests. The fragments with or without modifications are shown in red and black, respectively. Measurement was conducted in positive polarity mode.

### Supplementary Table 1. Prediction of E. coli tRNA modifications using tRNA-seq data

| Modification | Misincorporation | | | | Termination | | | | Total | tRNA species |
|---|---|---|---|---|---|---|---|---|---|---|
| | Frequency (%) | Position | Fraction >5%*[2] | Prediction*[3] | Frequency (%) | Position*[4] | Fraction >5%*[5] | Prediction*[7] | Prediction*[8] | |
| $s^{4}U$ | 4.5 - 50 | 8,9 | 28/29 | + | 0 - 42.9 | 10,11 | 9/29 | - | + | Ala1B, Arg2, Asn, Asp, Cys, Gln2, Gln1, Gly1, His, Ile2, fMet1, fMet2, Leu5, Met, Phe, Pro1, Ser3, Ser5, Ser1, Trp, Tyr1, Tyr2, Val2A, Val2B, Val1 |
| DD*[1] | 0 - 2.4 | 16-17, 20-20A | 0/20 | - | 0 - 51.5 | 17*[4], 20A*[4] | 18/20 | + | + | Gly3, His, Leu1A, Leu2, Lys, Met, Thr1, Thr2, Thr3, Trp, Val2A, Val2B, Asp, Ile1, Ile2, Ser1, Ser5 |
| Gm | 0 - 18.8 | 18 | 1/13 | - | 0 - 19.3 | 20 | 7/13 | + | + | Gln2, Gln1, Ile2, Leu5, Leu1A, Leu2, Leu3, Met, Ser2, Ser5, Ser1, Tyr1, Tyr2 |
| $s^{2}C$ | 1.7 - 7.3 | 32 | 2/4 | + | 1.5 - 26.4 | 34 | 3/4 | + | + | Arg2, Arg3, Arg4, Ser3 |
| Um | 0.3 - 0.5 | 32 | 0/4 | - | 0.7 - 10.7 | 34 | 2/4 | + | + | Pro1, Pro3, Gln2, Gln1 |
| Q | 0.8 - 11.6 | 34 | 1/4 | - | 4.5 - 62.9 | 36 | 3/4 | + | + | Tyr1, Tyr2, His, Asn |
| Glu-Q | 12.1 | 34 | 1/1 | + | 14.9 | 36 | 1/1 | + | + | Asp |
| $mnm^{5}s^{2}U$ | 4.6, 4.9 | 34 | 0/2 | - | 15.1, 19.9 | 36 | 2/2 | + | + | Glu, Lys |
| $cmnm^{5}s^{2}U$ | 6.5 | 34 | 1/1 | + | 5.3 | 36 | 1/1 | + | + | Gln1 |
| $cmnm^{5}Um$ | 11 | 34 | 1/1 | + | 62.6 | 36 | 1/1 | + | + | Leu4 |
| I | 99.9 | 34 | 1/1 | + | 0.5 | 36 | 0/1 | - | + | Arg2 |
| $k^{2}C$ | 91.7 | 34 | 1/1 | + | 96.1 | 35*[4] | 1/1 | + | + | Ile2 |
| $m^{1}G$ | 47.3 - 94.8 | 37 | 8/8 | + | 11.1 - 62.3 | 39 | 8/8 | + | + | Leu1A, Leu1B, Leu2, Leu3, Arg3, Pro1, Pro2, Pro3 |
| $ms^{2}i^{6}A$ | 5.6 - 56.2 | 37 | 9/9 | + | 71.8 - 91 | 39 | 9/9 | + | + | Leu5, Phe, Leu4, Trp, Cys, Ser2, Ser1, Tyr1, Tyr2 |
| $m^{6}t^{6}A$ | 1.9, 2.5 | 37 | 0/2 | - | 12.7, 13.1 | 37*[4] | 2/2 | + | + | Thr1, Thr3 |
| $acp^{3}U$ | 28.8 - 67.5 | 47 | 8/8 | + | 23.7 - 55.9 | 48*[4] | 8/8 | + | + | Phe, Val2A, Val2B, Arg2, Met, Ile2, Ile1, Lys |
| Ψ | 0 - 0.7 | 13,32,38, 39,40, 55,65 | 0/68 | - | 0 - 8.2 | 15,34,40, 41,42, 57,67 | 2/68 | - | - | Ala1B, Ala2, Arg2, Arg3, Arg4, Asn, Asp, Cys, fMet1, fMet2, Gln1, Gln2, Glu, Gly1, Gly2, Gly3, His, Ile1, Ile2, Leu1A, Leu2, Leu4, Leu5, Lys, Met, Phe, Pro1, Ser1, Ser2, Ser3, Ser5, Thr1, Thr2, Thr3, Trp, Tyr1, Tyr2, Val1, Val2A, Val2B |
| D"[1] | 0 - 0.9 | 16,17, 20,20A | 0/34 | - | 0 - 23.6 | 18,19, 21,22 | 3/34 | - | - | Leu5, Leu4, Phe, Trp, Gly1, Gly3, Leu1A, Leu2, Pro1, Gln2, Gln1, His, fMet1, fMet2, Thr2, Thr1, Thr3, Lys, Asn,Val1, Ala2, Ala1B, Arg3, Arg2, Ile1,Leu5, Asp,Ser2, Ser3 |
| Cm | 0 - 0.3 | 32 | 0/5 | - | 0.1 - 10.7 | 34 | 1/5 | - | - | Trp, Ser1, fMet1, fMet2, Thr4 |
| $ac^{4}C$ | 1.9 | 34 | 0/1 | - | 0.6 | 36 | 0/1 | - | - | Met |
| $cmo^{5}U$ | 0.3 | 34 | 0/2 | - | 0.9, 1.4 | 36 | 0/2 | - | - | Val1, Leu3 |
| $mcmo^{5}U$ | 0.1 - 0.5 | 34 | 0/4 | - | 1.2 - 8.1 | 36 | 1/4 | - | - | Ser1, Ala1B, Pro3, Thr4 |
| $mnm^{5}U$ | 0.2 | 34 | 0/2 | - | 0.3, 0.9 | 36 | 0/2 | - | - | Gly2, Arg4 |
| $m^{2}A$ | 0.2 - 1.5 | 37 | 0/6 | - | 0.2 - 0.9 | 39 | 0/6 | - | - | Asp, Glu, Gln2, Gln1, His, Arg2 |
| $m^{6}A$ | 0.2 | 37 | 0/1 | - | 0.2 | 39 | 0/1 | - | - | Val1 |
| $ct^{6}A$ | 0.3 - 2.9 | 37 | 0/8 | - | 0.4 - 5.4 | 39 | 1/8 | - | - | Met, Ile1, Ile2, Ser3, Arg4, Lys, Asn, Thr2 |
| $m^{7}G$ | 0.1 - 2.5 | 46 | 0/22 | - | 0.3 - 27.7 | 48 | 3/14*[6] | - | - | Ala2, Ala1B, Arg3, Arg2, Asn, Asp, Gly3, His, Ile1, Ile2, fMet1, Lys, Met, Phe, Pro1, Thr3, Thr1, Thr2, Trp, Val2B, Val2A, Val1 |
| T | 0 - 0.6 | 54 | 0/40 | - | 0 - 0.8 | 56 | 0/40 | - | - | Ala1B, Ala2, Arg2, Arg3, Arg4, Asn, Asp, Cys, fMet1, fMet2, Gln1, Gln2, Glu, Gly1, Gly2, Gly3, His, Ile1, Ile2, Leu1A, Leu2, Leu4, Leu5, Lys, Met, Phe, Pro1, Ser1, Ser2, Ser3, Ser5, Thr1, Thr2, Thr3, Trp, Tyr1, Tyr2, Val1, Val2A, Val2B |

*1. Note that single dihydrouridine (D) and tandem D lead to distinct RT signatures.

*2. The denominators used for these calculations are the number of modified sites established in the literature[3-10] and/or the database[2] (Supplementary Data 1). The numerators are the number of known modified sites where the misincorporation frequency was >5 % in tRNA-seq data. For example, 29 sites are reported to have $s^4U$ and the misincorporation frequency was > 5 % at 28 sites.

*3. Modifications were considered to be predictable (+) when the misincorporation fraction (previous column) was ≥ 50%.

*4. Positions of termination used for calculations were set empirically. Modifications led to a range of sites of termination. Most modifications led to the most pronounced termination 2 nucleotides downstream (3' side) from the modified site. For example, $m^1G$ or $ms^2i^6A$ at position 37 appear to lead to the termination of RT at position 39. However, at other sites, the highest termination occurs 1 nucleotide downstream from the modification; e.g., $acp^3U$ and $k^2C$ had peak termination at positions 48 and 35, respectively and tandem DD appears to induce termination at the latter D. In the case of $m^6t^6A$, termination appears at the same position as the modified site (position 37). The variation in the position of termination may be accounted for by variation in the mechanisms of inhibition of reverse transcription, e.g., based on the structure of modifications.

*5. The denominators used for these calculations are the number of modified sites established in the literature[3-10] and/or the database[2] (Supplementary Data 1). The numerators are the number of known modified sites where the termination frequency was >5 % in tRNA-seq data. For example, 4 sites are reported to have $s^2C$ and the termination frequency was > 5 % at 3 sites.

*6. Eight out of 22 tRNA species have both $m^7G$ and $acp^3U$ at position 46 and 47, respectively. Since $acp^3U$ has a strong effect on termination of RT, these sites were not considered in the assessment of $m^7G$'s effect on termination.

*7. Modifications were considered to be predictable (+) when the termination fraction (previous column) was ≥ 50%.

*8. A modification is considered predictable when the modification is predictable by misincorporation and/or termination.

**Supplementary Table 2. Strain list**

| **Strains** | **Relevant genotype/description** | **Reference/source** |
|---|---|---|
| *Vibrio cholerae* | C6706, wild-type El Tor clinical isolate ($Sm^R$) | 11 |
| *V. cholerae* Δ*thiI* | C6706 Δ*vc0894* | 1 |
| *V. cholerae* Δ*miaA* | C6706 Δ*vc0346* | 1 |
| *V. cholerae* Δ*ttcA* | C6706 Δ*vc1432* | This work |
| *V. cholerae* Δ*trmK* | C6706 Δ*vca0634* | This work |
| *V. cholerae* Δ*acpA* | C6706 Δ*vc0317* | This work |
| *V. cholerae vca0825*::*Tn* | C6706 *vca0825*::*Tn* | 12 |
| *V. cholerae vc1341*::*Tn* | C6706 *vc1341*::*Tn* | 12 |
| *V. cholerae vca0043*::*Tn* | C6706 *vc0043*::*Tn* | 12 |
| *V. cholerae vc0317*::*Tn* | C6706 *vc0317*::*Tn* | 12 |
| *V. cholerae vc0884*::*Tn* | C6706 *vc0884*::*Tn* | 12 |
| *Escherichia coli* MG1655 | wild-type MG1655 | |
| *Escherichia coli* DH5α λ*pir* | Cloning strain | |
| *Escherichia coli* SM10 λ*pir* | Conjugation donor | |
| *Vibrio parahaemolyticus* | RIMD 2210633 | 13 |
| *Aeromonas hydrophila* | ATCC7966 | ATCC |
| *Shewanella oneidensis* | MR-1 ATCC 700550 | ATCC |

**Supplementary Data Titles and Legends**

**Supplementary Data 1. Reference *E. coli* tRNA sequences with modifications.**
tRNA sequences are retrieved from tRNAdb[2] and partial or full sequences were changed or added based on the literature [3-10]. Bold letters indicate the changes and addition of sequences based on references.

**Supplementary Data 2. Conservation of tRNA modification enzymes between *V. cholerae* and *E. coli*.**

**Supplementary Data 3. Parameters of mass spectrometry for dynamic MRM analyses.**

**Supplementary Data 4. Comparative genomics for narrowing down candidate acetyltransferases required for acacp$^3$U biogenesis.**
Putative acetyltransferases in *V. cholerae* are listed with E-values calculated by BLAST among homologs between *V. cholerae* and indicated organisms. n.d. means the E-value is higher than 1E-10 or no detectable homologs were found.

**Supplementary Data 5. Primer list**

**Supplementary Data 6. Reference DNA sequences of tRNAs for mapping**
